# Polynomial Phylogenetic Analysis of Tree Shapes

**DOI:** 10.1101/2020.02.10.942367

**Authors:** Pengyu Liu, Priscila Biller, Matthew Gould, Caroline Colijn

## Abstract

Phylogenetic trees are a central tool in evolutionary biology. They demonstrate evolutionary patterns among species, genes, and with modern sequencing technologies, patterns of ancestry among sets of individuals. Phylogenetic trees usually consist of tree shapes, branch lengths and partial labels. Comparing tree shapes is a challenging aspect of comparing phylogenetic trees as there are few tools to describe tree shapes in a quantitative, accurate, comprehensive and easy-to-interpret way. Current methods to compare tree shapes are often based on scalar indices reflecting tree imbalance, and on frequencies of small subtrees. In this paper, we present tree comparisons and applications based on a polynomial that fully characterizes trees. Polynomials are important tools to describe discrete structures and have been used to study various objects including graphs and knots. There are also polynomials that describe rooted trees. We use tree-defining polynomials to compare tree shapes randomly generated by simulations and tree shapes reconstructed from data. Moreover, we show that the comparisons can be used to estimate parameters and to select the best-fit model that generates specific tree shapes.

---

A tree is a natural data structure that represents hierarchical relations between objects. In phylogenetics, a tree structure usually includes its tree shape, that is, the unlabeled underlying graph, as well as branch lengths reflecting either evolutionary distance or time. Estimating the branch lengths can be a challenge for tree reconstruction methods, with Bayesian and maximum likelihood methods yielding inconsistent results (Brown, 2010), high demands on memory and processor time (Binet, 2016), and/or lack of strong support for a molecular clock (in the case of timed trees). As a consequence, the inferred phylogenetic trees may have a consistent tree shape but differing root heights and branch lengths.

The shapes of phylogenetic trees can carry information about macroevolutionary processes, as well as reflecting the data used and the choice of the evolutionary model (Kirkpatrick, 1993; Purvis, 2011; Aldous, 1996). The ecological fitness and the presence of selection can also affect the shapes of trees (Dayarian, 2014; Maia, 2004). In the study of infectious diseases, where the shapes of phylogenetic trees of pathogens reveal diversity patterns that represent a combination of unfixed neutral variation, variation under selection, demographic processes and ecological interactions, it is not clear how informative the tree shapes are of the underlying evolutionary and epidemiological processes. However, effort is being made to explore this question, with the main focus often on the frequency of cherries and tree imbalance (Grenfell, 2004; Lambert, 2013; Plazzotta, 2016; Volz, 2013).

One of the main topics of inquiry in phylogenetic tree shapes has been asymmetry, since a key observation was made that the shapes of phylogenetic trees reconstructed from data are more asymmetric than tree shapes simulated by simple models (Aldous, 1996). Various ways to measure the asymmetry were developed (Aldous, 1996; Colless, 1982; Fusco, 1995; Sackin, 1972; Stich, 2009) and it was shown that these asymmetric measures can distinguish random trees generated by different models (Agapow, 2002; Kirkpatrick, 1993; Matsen, 2006). At the same time, mathematical models that produce imbalanced trees were developed (Aldous, 2001; Blum, 2006). As statistical tools, the distributions of tree shapes under simple models can be used to test evolutionary hypotheses (Blum, 2006; Mooers, 1997; Wu, 2016). In (Manceau, 2015), and mathematical models can be developed to match the macroevolutionary patterns observed in the phylogenetic trees reconstructed from data.

As the cost of DNA sequencing is decreasing, more genomic data are being collected and becoming available. More organisms are being sequenced progressively at the whole-genome scale (Bedford, 2015; Chewapreecha, 2014; Colijn, 2018) and the evolution of certain pathogens is being tracked in real time (Hadfield, 2018). As a consequence, both the number and the size of trees reconstructed from data are increasing. Accordingly, a major challenge in tree shape analysis is that there are few tools to describe and compare trees in a quantitative, accurate, comprehensive and easy-to-interpret way, especially for large trees. Scalar indices describing asymmetry or the frequency of subtrees have a limitation in that many different tree shapes may have the same index. A labelled tree is a tree shape whose vertices have unique labels. An alternative approach to comparing tree shapes is using metrics defined for labelled trees, for example, the well known Robinson-Foulds metric (Robinson, 1981), Billera-Holmes-Vogtmann metric (Billera, 2001) and Kendall-Colijn metric (Kendall, 2016), among others. These metrics depend on the labels of the vertices, that is, two labelled trees with the same tree shape but the labels re-arranged are not identical and the distances between them can be very large. Recently, metrics defined for rooted unlabelled trees or rooted tree shapes have also been introduced (Colijn, 2018), making use of integer labels assigned to tree shapes. However, these metrics have several limitations, including the challenge of interpreting the integer labels, the treatment of non-binary trees, and the metrics’ performance in distinguishing trees from different processes or datasets.

Graph polynomials and knot polynomials are important tools in the mathematical study of discrete structures, and can be used to describe the structures in interpretable ways. For example, the Tutte polynomial (Tutte, 1954) is a renowned polynomial for graphs and the Jones polynomial (Jones, 1985) is one of the most important tools to study knots. In (Liu, 2021), a method to assign a unique polynomial to each tree shape is introduced. These polynomials provide a new way to describe tree shapes quantitatively and comprehensively. The coefficients of the polynomial of a tree can be considered as a generalization of the clade size distribution of the tree. In addition, the set of coefficients of a tree polynomial can be treated as a vector, and vectors are natural objects on which to define metrics. In this paper, we introduce the polynomial representations for tree shapes and we define and examine a metric based on the trees’ unique polynomials. We show that the polynomial representations for tree shapes have perfect resolution and reasonably low computation time, and the polynomial metric has a performs well at clustering trees, compared to other high-resolution metrics. We also show that the polynomials can be used for parameter estimation, and for choosing the best-fit model to generate a tree shape.

## Materials and Methods

### Tree Polynomials

In this paper, a tree shape or simply a tree represents an unlabeled tree, that is a graph with no cycles, without information about branch lengths or labels unless otherwise stated. We define the bivariate polynomial *P*(*T, x, y*) for a rooted unlabeled tree *T* in the following way. If *T* is the trivial tree with a single vertex, then *P*(*T, x, y*) = *x*. Otherwise *T* has *k* branches at its root and each branch leads to a subtree of *T*. Let *T*_1_, *T*_2_, …, *T*_*k*_ be the *k* rooted subtrees whose roots are adjacent to the root of *T*. We define the polynomial for *T* by 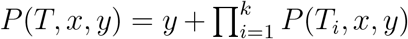. If all of the subtrees are the trivial tree, then the polynomial is defined and we have a rooted *k*-star whose polynomial is *P*(*T, x, y*) = *x*^*k*^ + *y*. If there exists a non-trivial subtree *T_i_*, then we apply the definition to compute *P*(*T*_*i*_, *x, y*). The polynomial *P*(*T, x, y*) can be computed by recursively applying the definition until we reach all tips of *T*. As another example, the polynomial for the three-tip rooted binary tree *T* is *P*(*T, x, y*) = *x*(*x*^2^ + *y*) + *y* = *x*^3^ + *xy* + *y*, as *T* has two subtrees adjacent to the root, a trivial tree *T*_1_ with *P*(*T*_1_, *x, y*) = *x* and a cherry *T*_2_ with *P*(*T*_2_, *x, y*) = *x*^2^ + *y*. It is proved in (Liu, 2021) that the polynomial distinguishes unlabeled rooted trees and can be generalized to distinguish unlabeled unrooted trees. A rooted tree can be reconstructed from its polynomial by computing its Newick code, which can be obtained by recursively subtracting *y* and factoring the rest of the polynomial. Methods to factor large multivariate polynomials can be found in (Monagan, 2018). The coefficients of a tree polynomial can be written as a matrix. Let *T* be a rooted tree with *n* tips. Its coefficient matrix *C*(*T*) or (*c*^(*a,b*)^) is displayed as follows, where *c*^(*a,b*)^ is the coefficient in the term *c*^(*a,b*)^*x*^*a*^*y*^*b*^.

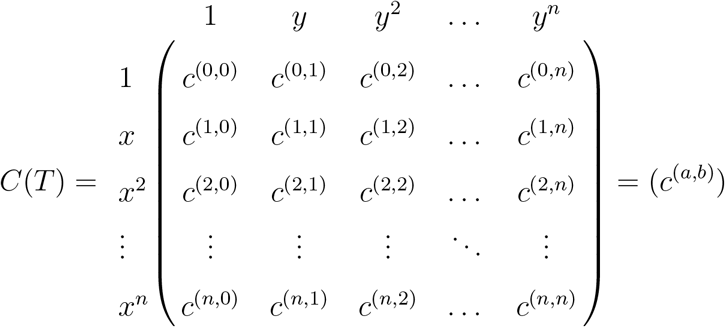

Let *T* be a rooted tree with *n* tips. The coefficient *c*^(*a,b*)^ in the term *c*^(*a,b*)^*x*^*a*^*y*^*b*^ of *C*(*T*) can be interpreted as the number of ways in *T* to choose *b* clades (with more than one tip) such that these clades include *n − a* tips of *T* in total. The clade size distribution of a tree *T* is the vector whose *i*-th element is the number of clades in *T* containing *i* tips. The second column in the matrix *C*(*T*) is the clade size distribution of the tree *T*, where *c*^(*n−k*,1)^ indicates the number of clades with *k* tips (Liu, 2021). It is also showed in (Liu, 2021) that if we substitute the variable *y* in a polynomial *P*(*T, x, y*) by a prime number or a Gaussian prime *p*, the resulting polynomial *P*(*T, x, p*) can still distinguish all rooted binary trees. This property of the polynomial can be utilized to make tree analysis faster.

### Tree metrics

In this paper, we use three tree metrics or distances. The first is a tree metric based on the Laplacian spectrum. The metric is the Jensen-Shannon distance over the spectrum densities introduced in (Lewitus, 2016). We call it Lewitus-Morlon metric. The second metric is based on the subtree size distribution. The subtree size distribution of a tree is defined as a vector whose *n*-th entry is the number of *n*-tip subtrees in the tree. The metric is defined using the Manhattan distance over the subtree size distribution vectors. We name it the “subtree-Manhatttan metric”. The third metric is based on the polynomial. Let *T*_1_, *T*_2_ be two trees and 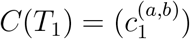, 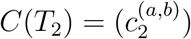 be the coefficient matrices of the polynomials *P*(*T*_1_, *x, y*), *P*(*T*_2_, *x, y*). We define a function

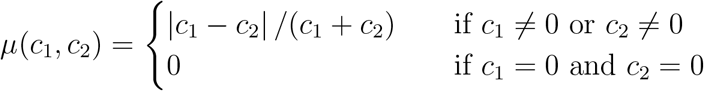

and the metric by

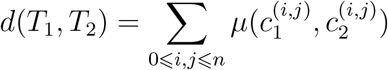

This metric is not only defined for trees of the same size, but also for trees of different sizes where it’s natural to assign a coefficient of 0 to each term that is absent in a polynomial.

### Parameter estimation and model selection

To estimate parameters for trees, we use the polynomial metric or the subtree-Manhattan metric together with the weighted average of the neighboring observed data with the nearest neighbor kernel smoother. Specifically, we generate a set of observed trees 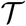 using a random tree generator with the different vectors of parameters *ρ*. For any tree *T* in 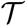, let *ρ*(*T*) be the vector of parameters used to generate *T*. We estimate the parameters of a tree *T*_0_ by the weighted average as follows:

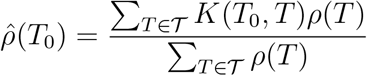

where *K*(*T*_0_, *T*) is the *k*-nearest-neighbor kernel function, that is, *K*(*T*_0_, *T*) = 1*/k* if *T* is a *k* nearest neighbor of *T*_0_ under the polynomial metric and *K*(*T*_0_, *T*) = 0 otherwise. We choose different *k* for different sets of observed trees. For a set of observed trees 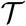, we generate another set 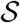 of 1,000 random trees. For each *k* from 1 to 20, we estimate parameters of trees in 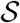 using the set of observed trees 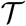, and we have the average estimation error for each *k*. We choose the *k* that has minimum average estimation error for the set of observed trees 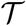.

We use naive Bayes classifiers (Rish, 2001) together with the polynomial to perform model selection. Naive Bayes classifiers assume independence of the predictor variable. We label each tree according to the underlying model (beta splitting, the explosive radiation and trait evolution), and use the trees’ polynomial coefficients as features.

### Simulations

#### Beta splitting trees

The beta splitting random trees used in this paper are generated by the beta-splitting model introduced in (Aldous, 1996). At each branching event, the probability of one child clade containing *i* tips and the other child clade containing *n* − *i* tips is given by the following formula.

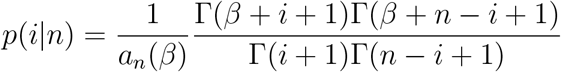

The Γ(*z*) in the formula is the Gamma function and *a*_*n*_(*β*) is a normalizing constant.

Our sets of *n*-tip modeled beta splitting trees consist of trees generated with *β* = 0, *β* = −1, and *β* = −1.5, and there are 100 trees for each parameter. These choices of *β* correspond to the Yule model, the Aldous branching model and the proportional to distinguishable arrangements (PDA) model (Blum, 2006). We also use sets of beta splitting trees consisting of 1,000 such trees, with *n* tips and parameters *β* that are uniformly randomly chosen from the interval [−1.5, 8.5].

#### Explosive radiation trees

The explosive radiation trees were simulated with a modification of the birth-death model proposed by Steel (2001). Steel’s model builds on the traditional constant birth-death model by setting lineage-specific speciation rates. More precisely, the rate of speciation events on a given lineage is a function of *t*, the time to the last speciation event on that lineage. This time *t* is reset to 0 at every speciation, and the birth (λ_*i*_) and death (*μ*_*i*_) rates of a given lineage *i* are then defined as follows:

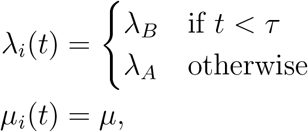

where λ_*A*_, λ_*B*_, *μ* and *τ* are parameters of the model.

All rates are defined as the number of events per tip per time unit. The choice of the time unit is not relevant to our experiments, as the polynomial does not make use of information on branch lengths.

A data set of *n*-tip explosive radiation trees contains 1,000 random trees generated with the birth rate λ_*B*_ fixed at 1.0 (per arbitrary time unit), the time shifting the birth rates *τ* fixed at 0.5 time unit, and both the birth rate λ_*A*_ and the death rate *μ* uniformly randomly chosen from the interval [0, 1].

#### Trait evolution trees

This data set was simulated following the birth-death model proposed by Heard (1996). In this model, each lineage has an associated trait value (*x*) which is “inherited” at speciation events with some stochastic change. The model for trait evolution implemented here is a linear-Brownian variation, where additive changes are made to the trait value at each speciation event: *x*_new_ = max{*x*_old_ + *ϵ*, 0.01}. The stochastic change *ϵ* is drawn from a normal distribution with expectation zero and standard deviation *σ*_*x*_. Both *σ*_*x*_ and *x*_0_ (the trait value at the root) are parameters of the model.

The birth (λ_*i*_) and death (*μ*_*i*_) rates are defined as λ_*i*_ = *x* and *μ*_*i*_ = *μ*, respectively. Similar to the explosive radiation model, the death rate *μ* is constant in time and across lineages. Notice that there are numerous ways to produce trees with a given number of species from an evolutionary model (Hartmann, 2010). For all evolutionary models used in our analysis, trees are simulated forward in time until *n* tips are first reached. Our data sets of *n*-tip trait evolution trees contain 1,000 random trees generated with the initial birth rate fixed at 1.0 (per arbitrary time unit), and the birth rate variation at a speciation event and the death rate uniformly randomly chosen from the interval [0, 1].

We do not down-sample the simulated trees despite the fact that the data we use (see below) are only a small minority of the true numbers of tips in the relevant settings. This would be infeasible at genuine scales given the comparatively high true population sizes of circulating pathogens. For example, only a very small minority of circulating influenza infections lead to a sequence deposition in the database, with many others going undetected and/or unsequenced. Those that are sequenced may not be unique exemplars of their sequences in the population, as transmission may occur without detectable variation. As a consequence, in comparing simulation models to data, we interpret simulated branching events as diversification events that are likely to be ancestral to sampled tips and therefore observed, and “death” events as, effectively, sampling events that stop onward transmission of the particular lineage.

### Data

#### HIV and influenza virus trees

The HIV trees were described and analyzed previously (Chindelevitch, 2019). Briefly, HIV-1 sequence data from three studies were used. The Wolf et al. study (Wolf, 2017) provided data from a concentrated epidemic of HIV-1 subtype B, occurring primarily in men who have sex with men (MSM) in Seattle, USA. The Novitsky et al. study (Novitsky, 2013) describes data from a generalized epidemic of HIV-1 subtype C in Mochudi, Botswana, a village in which the HIV-1 prevalence in the adult population at the time was estimated to be approximately 20%. Hunt et al. (Hunt, 2013) describes data from a national survey of the generalized epidemic of HIV-1 subtype C in South Africa. These datasets reflect a diverse set of spatial scales and epidemiological contexts. Phylogenetic reconstruction was described in (Chindelevitch, 2019); briefly, trees were reconstructed using RAxML (Stamatakis, 2014), which is a maximum likelihood method, under a general time-reversible (GTR) model of nucleotide substitution. We use a GTRCAT model for rate variation among sites. Each tree was based on a random sample of 100 sequences. We use a subtype D sequence as an outgroup to root HIV-1 subtype B phylogenies.

Our influenza virus trees were previously described in (Colijn, 2018). We aligned HA protein sequences from NCBI, focusing on human influenza A (H3N2). Data were downloaded from NCBI on 22 Jan. 2016. We included full-length HA sequences with collection date. The USA dataset (*n* = 2168) includes sequences from the USA with collection dates between Mar. 2010 and Sep. 2015. The tropical dataset (*n* = 1388) includes sequences with a location listed as tropical, with collection dates within Jan. 2000 and Oct. 2015. Accession numbers are included in the Supporting Information of Colijn (2018). Fasta files were aligned with mafft, and for both the tropical and USA datasets, 500 taxa were selected uniformly at random 200 times. We inferred 200 corresponding phylogenetic trees with FastTree (Price, 2010). Where necessary we re-aligned the 500 selected sequences before performing tree inference. This process resulted in 200 “tropical” influenza virus trees and 200 “USA” influenza virus trees, each with 500 tips, reconstructed from the HA region of human H3N2 samples. Note that this approach is distinct from the perhaps more familiar phylogenetic methods where bootstrapping or Bayesian reconstructions results in many trees on *one* set of tips. These are likely to share features and structures because they describe the ancestry of the same set of taxa. Here, each tree has a different set of tips (though there is some overlap).

#### WHO influenza virus clades

We used several influenza virus clades, described in (Hayati, 2020). In that work we downloaded all human H3N2 full-length HA sequences with dates between 1980 and May 2018 and created a large, timed phylogeny of H3N2 using RAxML and Least Squares Dating (Stamatakis, 2014; To, 2016). This “full” tree has over 12,000 tips. We used the Nextflu (Neher, 2015) *augur* pipeline (https://bedford.io/projects/nextflu/augur/) to assign a WHO clade designation to the sequences. The WHO defines named clades using specific mutations in the HA1 and HA2 subunits of the HA protein. The full list of mutations is available at: https://github.com/nextstrain/seasonal-flu/blob/master/config/clades_h3n2_ha.tsv. We assign a sequence to a clade if it contains all the mutations defining that clade. We then extracted the subtrees of the “full” tree corresponding to specific WHO clades A1B/135N (60 tips), A1B/135K (63 tips), 3c3.B (117 tips) and A3 (227 tips). These are recent and appropriately-sized trees which we use here to demonstrate parameter estimation for simple models, and model selection among our four random tree models.

**Table 1.**
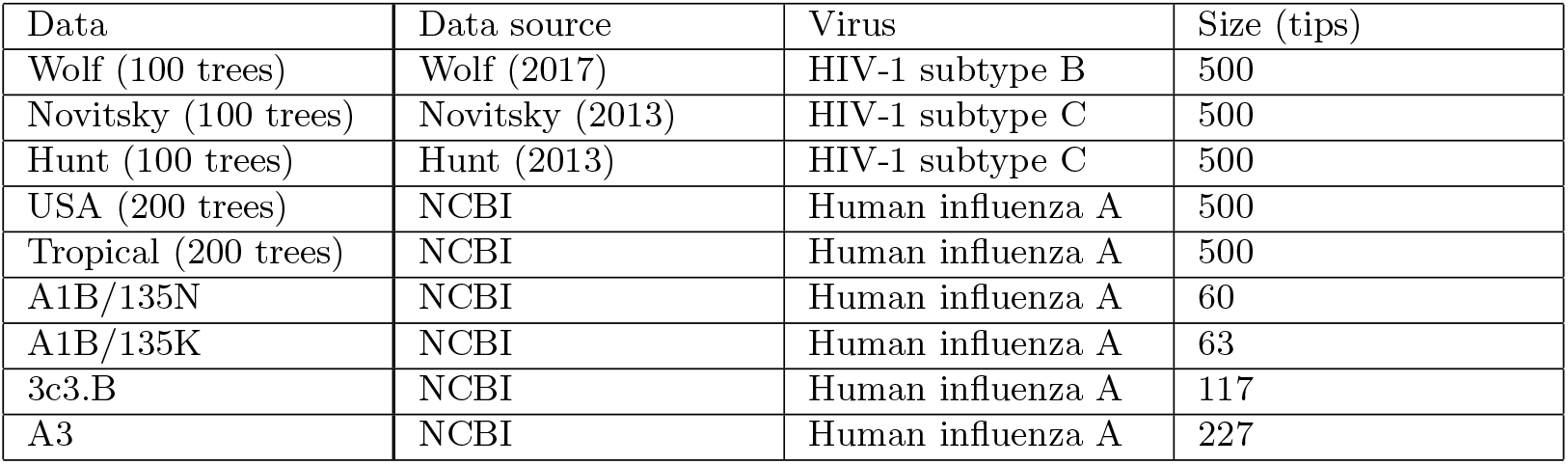
Summary of virus phylogenies.

### Implementation

We developed an R package named *treenomial*, which is available at CRAN. We also prepared a demonstration named *treeverse*, which displays a 3-dimensional projection of the polynomial metric space of all binary tree shapes up to 16 tips with interactive options available at https://magpiegroup.shinyapps.io/treeverse/.

## Results

### Tree Representations and Metrics

We compare the polynomial to other tree representation methods in terms of computation time and resolution. These tree-representing methods include the Colless index (Agapow, 2002), gamma statistics (Pybus, 2000), the Sackin index (Sackin, 1972), the subtree size distribution and more recently introduced Laplacian spectrum (Lewitus, 2016). The resolution of a tree-representing method (for *n*-tip trees) is defined to be the ratio of the number of unique representations to the total number of non-isomorphic tree shapes with *n* tips. We compute these representations for all tree shapes with 15 tips (where there are 4850 non-isomorphic tree shapes). Figure 1 A displays the computation time and the resolution of these methods, where the data point “combined” is the vector comprising the Colless index, gamma statistics and the Sackin index. The results show that Laplacian spectrum, the polynomial, and the subtree size distribution (with more than one parameter) have higher resolution than scalar summary statistics while the scalar Colless index, gamma statistics and the Sackin index have lower resolution. As there are vastly numerous non-isomorphic tree shapes with hundreds of tips, it is not feasible to compute the resolution for larger trees, but we know that the resolution of the subtree size distribution decreases as the number of tips increases, and the Laplacian spectrum is not guaranteed to have 100% resolution for all trees, that is, there are non-isomorphic trees with the same spectrum density (Lewitus, 2016). The polynomial, on the other hand, is guaranteed to distinguish all trees (Liu, 2021). In Figure 1 B, we show how computation time of the subtree size distribution, the Laplacian spectrum and the polynomial for a single tree changes as the size of trees increase. Among the high-resolution tree-representing methods we compared, the polynomial has low computation time and keeps the resolution at 100% for trees of any size.

**Figure 1.**
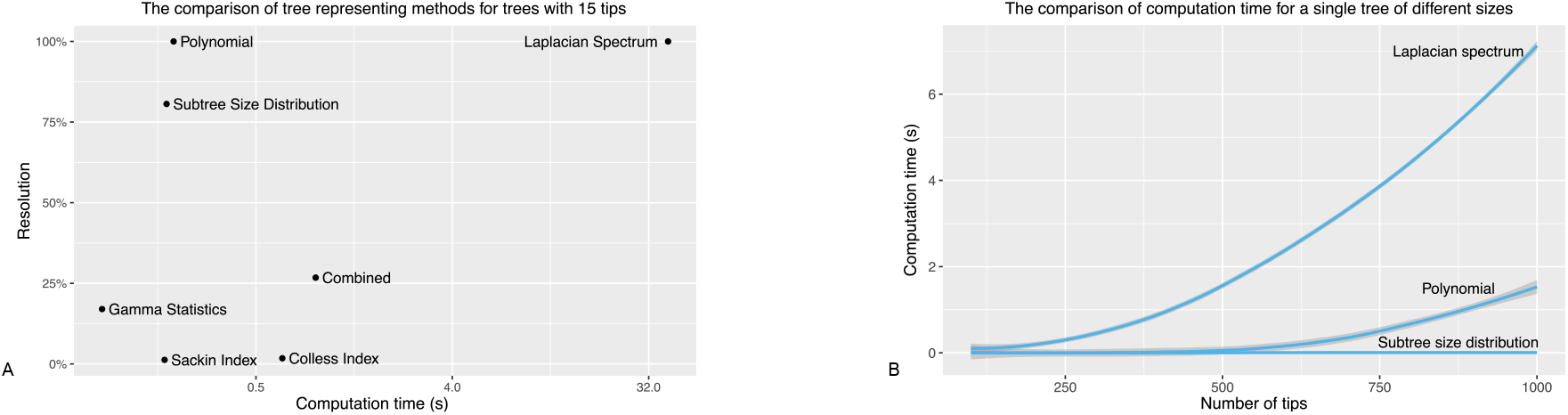
A: the comparison of tree representing methods, where the combined is the a combined vector of the Colless index, gamma statistics and the Sackin index. B: the comparison of the computation time for trees of different sizes. These are based on the computation time for random trees with 100 to 1,000 tips with increment of 50 tips; each data point denotes the average computation time over 1,000 trees.

Tree representations can induce tree metrics, which are important tools in comparing phylogenetic trees. We compare the polynomial metric with the metric induced by high-resolution tree-representing methods, that is, Lewitus-Morlon metric and the subtree-Manhattan metric. The polynomial metric is a genuine metric on trees, in the sense that it only gives a distance of zero if two trees have identical shapes, it is symmetric, and obeys the triangle inequality (see the supplement for proof; in contrast, the subtree-Manhattan metric and Lewitus-Morlon metric are not metrics in the mathematical distance sense (Lewitus, 2016)). The polynomial metric also has the advantage that the distance between a pair of trees is bounded above by the number of non-zero entries in the coefficient matrix of the larger tree. More precisely, let the larger tree be of *n* tips; the polynomial distance between the trees has an upper bound of *n*⌊*n*/2⌋ − ⌊*n*/2⌋^2^. The distribution of the pairwise distances between trees of the same size resembles a normal distribution, which gives a relative reference for how large the distance between a pair of trees is compared to what one might expect. See Supplementary Figure 1 for the distribution. Figure 2 A-C displays visualizations of the three distances between trees in a data set of 100-tip modeled beta splitting trees. We apply the *k*-medoids clustering algorithm PAM described in (Kauffman, 1990) to, respectively, the Lewitus-Morlon distance matrix, the subtree-Manhattan distance matrix and the polynomial distance matrix of a set of the 100-tip modeled beta splitting trees. We repeat this experiment for 100 times; Figure 2 D shows the misclassification rates. The polynomial metric has smaller misclassification rates than the other two metrics, which indicates that the polynomial has the potential to perform better in tasks involving clustering phylogenetic trees.

**Figure 2.**
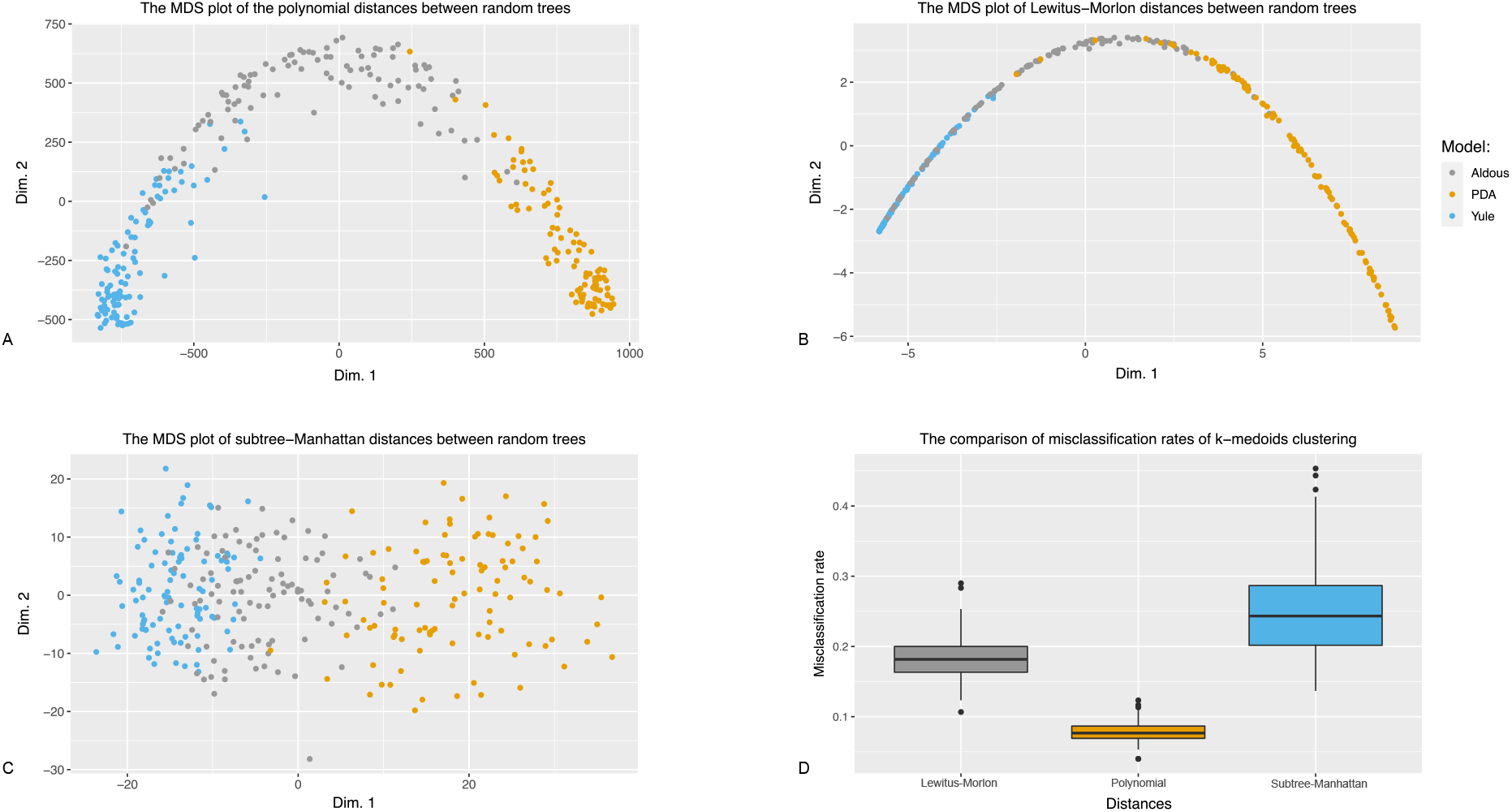
A-C: the multidimensional scaling plots of the three distances between trees in a set of 100-tip random trees, where each dot represent a random tree. D: the comparison of the misclassification rates of *k*-medoids clustering.

### Parameter Estimation and Model Selection

#### Parameter estimation

We show that the polynomial can be used to create likelihood free methods for parameter estimation. Here, we display the results of parameter estimation using the polynomial metric together with a simple weighted average method described in the method section. We generate a set of 250-tip beta splitting trees and use the set of random trees as observed data in the parameter estimation method; we then estimate the parameter *β* for 100 beta splitting trees with 250 tips. Figure 3 A shows the result of the estimation, and Figure 3 B shows the result of the estimation for 500-tip beta splitting trees. See Supplementary Table 1 for the summary of the estimation. In general the estimation works better for larger trees, and is better when the parameter *β* is in the interval [−1.5, 2]. We note that where a likelihood model is available, maximizing the likelihood may well be better than likelihood-free inference based on tree descriptions, but these results indicate that the polynomial contains relevant information that high-performance likelihood-free inference methods could utilize.

**Figure 3.**
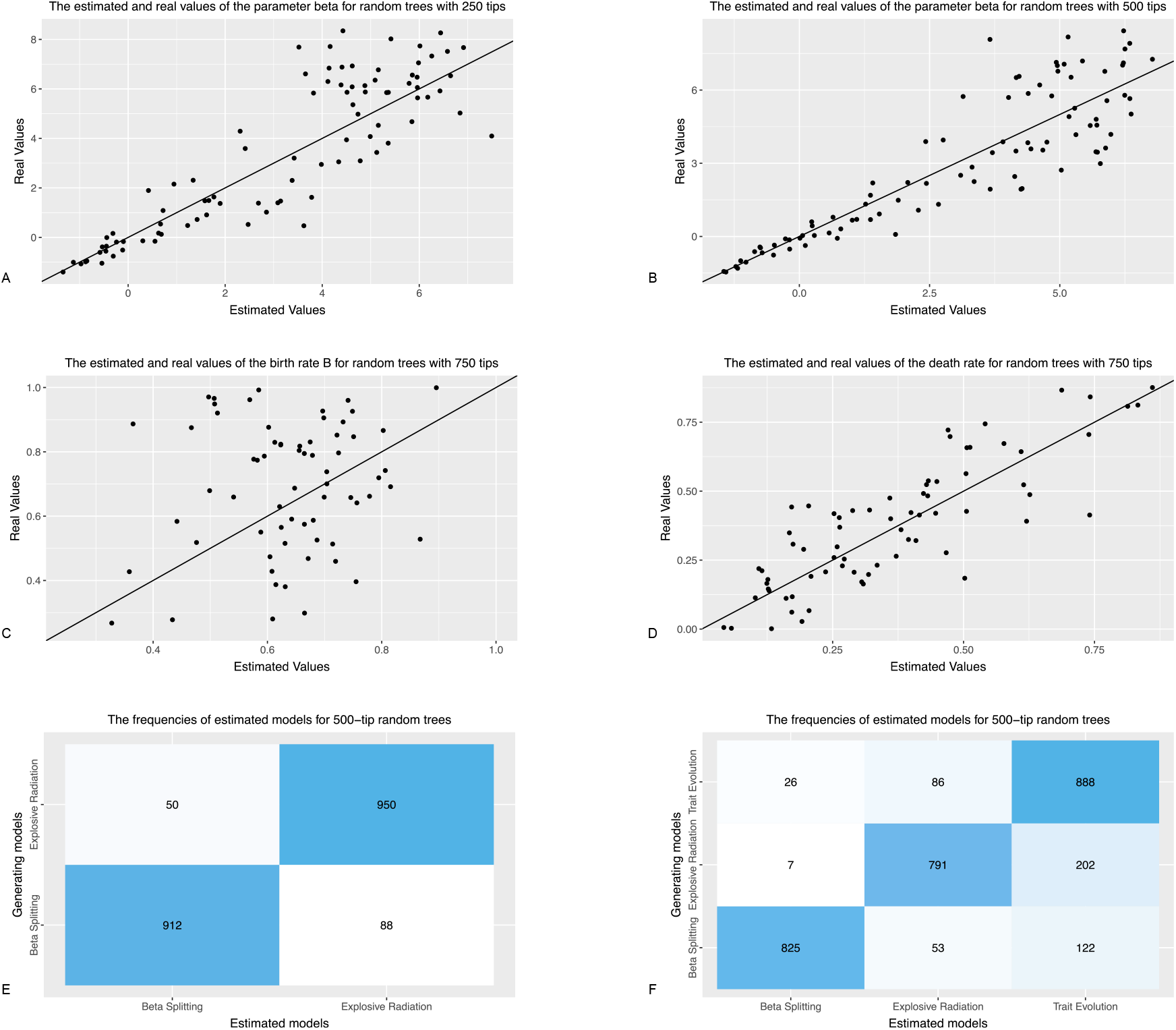
A-B: the comparisons between the real parameter and the estimated parameter of the beta splitting random trees with 250 tips and 500 tips using polynomials. C-D: the comparisons between the real parameters and the estimated parameters of the explosive radiation random trees with 750 tips using polynomials. E-F: the results of using naive Bayes classifiers to select the model generating random trees with 500 tips using polynomials.

We also generate a set of 750-tip explosive radiation trees and use the set of random trees as observed data in the parameter estimation method to estimate the birth rate λ_*B*_ and death rate *μ* for 100 explosive radiation trees with 750 tips. Figure 3 C-D shows the results of the estimation. The results are not as good as the results for beta splitting trees, especially the results for the birth rate λ_*B*_. Supplementary Tables 1–3 give details of the relationship between estimated and true values. We also use the subtree-Manhattan metric and the same weighted average method to perform parameter estimation for the same data sets. See Supplementary Figure 4; we find that the polynomial metric performs better than the subtree-Manhattan metric with the weighted average method in estimating parameters for both beta splitting trees and the explosive radiation trees.

#### Model selection

The beta splitting model and the explosive radiation model are different random tree generators. The beta splitting model uses the Markov branching process while the explosive radiation model uses a birth-death process. Both processes are commonly used in random tree generators, for example, the trait evolution model is another tree generator based on the birth-death process. Figure 2 shows that the polynomial has the potential to distinguish different tree generating models. In this section, we use the polynomial together with naive Bayes classifiers to estimate which model is used to generate a tree.

We generate a set of 500-tip beta splitting trees, a set of 500-tip explosive radiation trees, and a set of 500-tip trait evolution trees. We use these sets of random trees as observed data together with the naive Bayes classifiers to classify random trees generated by these three models. Figure 3 E shows the results of the experiment where we only use the set of beta splitting trees and the set of explosive radiation trees to train the naïve Bayes classifier, and use the classifier to classify a set of 1,000 beta splitting trees and 1,000 explosive radiation trees. Figure 3 F shows the results of the experiment where we use all three sets of random trees to train the naive Bayes classifier and use the classifier to classify a set of 1,000 beta splitting trees, 1,000 explosive radiation trees, and 1,000 trait evolution trees. The accuracy of the first experiment is 93.1% and of the second experiment is 83.5%, where the main misclassification (58.1% of the misclassified cases) is between the explosive radiation model and the trait evolution model, the two models based on the birth-death processes. Supplementary Figure 3 shows the results for 250-tip and 750-tip trees, and that this model selection method is more robust with larger trees. The results show that the polynomial together with naive Bayes classifiers can be a good tool in finding a tree generator that fits a given tree, as not only are trees from different random processes distinguished well, but the two different birth-death processes are also well distinguished.

We also use the subtree size distribution and the standard naive Bayes classifiers to perform model selection for the same data sets. See Supplementary Figure 4. Compared to the polynomial, the accuracy of the first experiment using the subtree size distribution is 82.7% and of the second experiment is 71.6%, where more explosive radiation trees are classified as beta splitting trees. To further understand the differences between the polynomial and the subtree size distribution in the naive Bayes classifiers, we display the most informative features in the classifiers in Supplementary Figure 5. It shows that for the subtree size distribution, the most informative features in model selection are the number of subtrees with approximately 400 tips, which could be considered as a description of tree imbalance for more imbalanced trees would have more subtrees with 400 tips than the balanced ones. On the other hand, Supplementary Figure 5 B shows that other than the clade size distribution (the dark thin strip at the bottom), the most informative features for the polynomial also include the coefficients in the black area at the top, which are interpreted as the numbers of ways to choose as many clades (with more than one tips) as possible so that the clades contain all or most of the tips in a tree. Compared to the subtree size distribution, this extra information gives the performance of the model selection method a boost.

### Applications to Data

Human influenza virus A H3N2. Influenza virus A is highly seasonal outside the tropics and most cases occur in the winter (Russell, 2008), whereas there is relatively little seasonal variation in the tropics. This demonstrative data set provides trees for the same virus circulating with different epidemiological dynamics (seasonal forcing in temperate regions, vs lack of seasonality in the tropics). The second data set consists of three samples of trees inferred from HIV-1 sequences in different settings: subtype B among men who have sex with men (MSM) in Seattle (Wolf, 2017), HIV 1C circulating at the village scale in Botswana (Novitsky, 2013) and a national-level dataset from South Africa (Hunt, 2013). As with influenza virus, it is to be hoped that these different epidemiological patterns are revealed in the shapes of reconstructed phylogenetic trees (Chindelevitch, 2019; Colijn, 2018).

We visualize the polynomial distances between trees in these two sets by classical MDS in Figure 4 A-B. We also use the *k*-medoids clustering on the data and we have the results displayed in Figure 4 C-D. The influenza trees are very well separated into desired groups under the *k*-medoids clustering. This indicates that classifying the epidemiological process behind a tree using the polynomial metric would likely be possible. In the supplement, we also compute the binary differences (Choi, 2010) of the polynomials for these trees, which improves the results of the *k*-medoids clustering. See Supplementary Figure 6. For these particular challenges, however, typically a researcher would know whether their data were from the tropics or not, or what the broad epidemiological setting (village, national, Western population MSM) was at the time of collection. We therefore focus on more specific estimation questions (parameter estimation and model choice).

**Figure 4.**
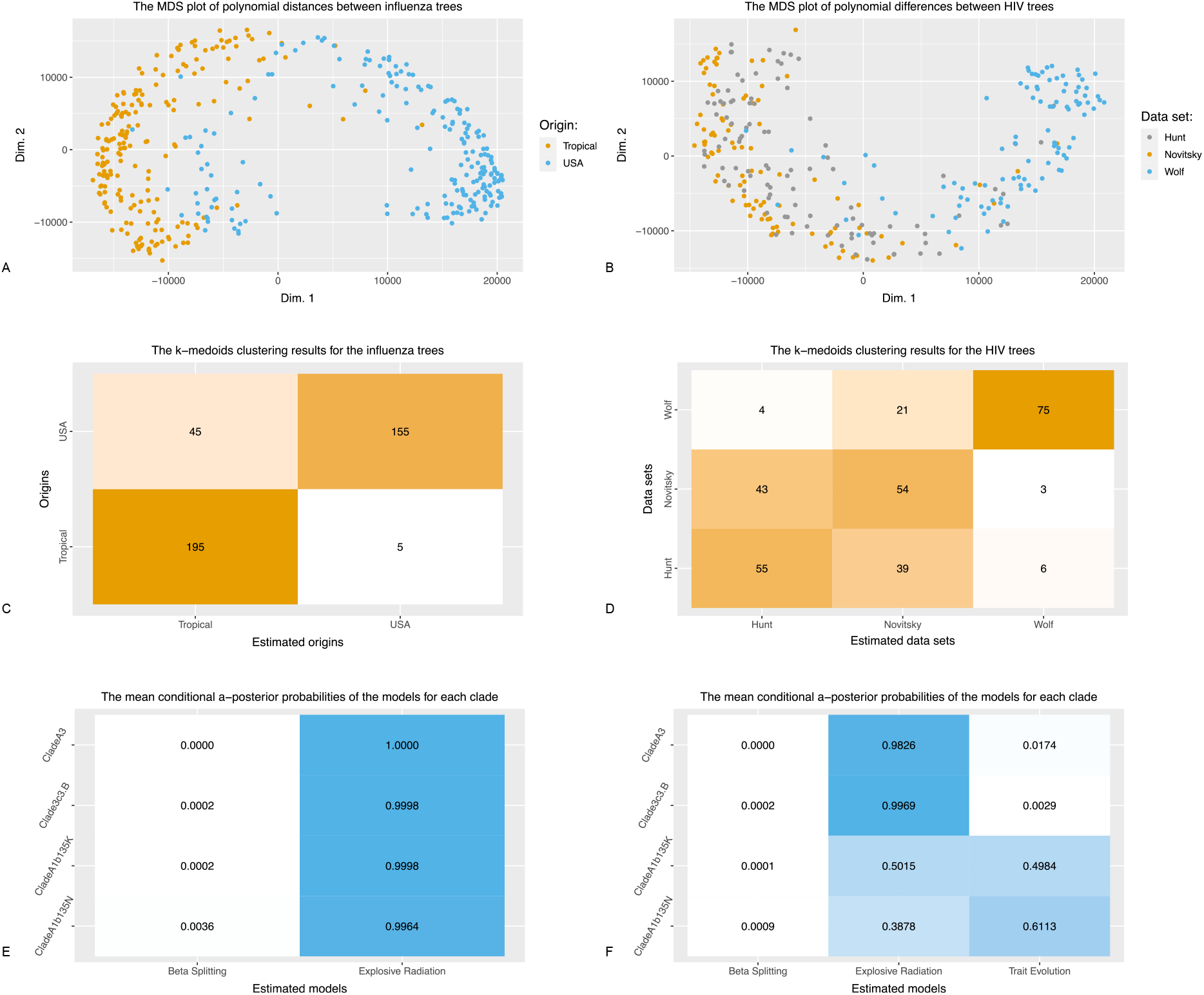
A: the MDS plots of the polynomial distances between the influenza trees. B: the MDS plots of the polynomial distances between the HIV trees. C: the results of *k*-medoids clustering for the influenza trees using the polynomial metric. D: the results of *k*-medoids clustering for the HIV trees using the polynomial metric. E-F: the mean conditional a posteriori probabilities (over the 1,000 naive Bayes classifiers) of the model estimation for the influenza clades.

As an example of applying the parameter estimation and model selection methods to data, we first select the models that best fit the four WHO influenza clades, A1B/135N (60 tips), A1B/135K (63 tips), 3c3.B (117 tips) and A3 (227 tips), then estimate the parameters for the model that best fits the clade. To select the model that best fits a clade, we generate a set of beta splitting trees, a set of of explosive radiation trees and a set of trait evolution trees of the same size as the clade. We use these sets of trees and naive Bayes classifiers to estimate the a-posterior probabilities of the clade being generated by the models. Figure 4 E shows that if we select only from the beta splitting model and the explosive radiation model, then all four clades are deemed more likely to be generated by the explosive radiation model, a tree generator based on the birth-death model. Figure 4 F shows that if we include the trait evolution model, the small clades A1B/135N (60 tips) and A1B/135K (63 tips) are predicted to be generated by either the explosive radiation model or the trait evolution model. The classifiers predict that for larger clades, the most likely model is the explosive radiation model. Both models seem reasonable for influenza, as a new variant that has polymorphisms allowing it to evade immunity that has built up in the population due to exposure to previous influenza viruses could have an early rapid rise (explosive radiation). However, influenza viruses have numerous traits (including interactions with host immunity) that could influence the branching rates in influenza virus phylogenies.

As an example, we examine influenza virus clade A3 (227 tips) in detail and estimate its parameters. First, we generate a set of beta splitting trees, a set of explosive radiation trees, and a set of trait evolution trees, all with 227 tips. For each set, we choose 250 random trees to visualize. In total we thus have 751 trees including clade A3. Figure 5A shows a visualization of the polynomial distances among these trees. Figure 5B shows the results of estimating the parameters (repeated 100 times with different sets of random trees) of the explosive radiation model for clade A3. The 95% confidence interval of the birth rate λ_*B*_ is (0.50, 0.54) and the 95% confidence interval of the death rate *μ* is (0.52, 0.56). The 95% confidence interval of *R*_0_ of the clade is (0.906, 1.019).

**Figure 5.**
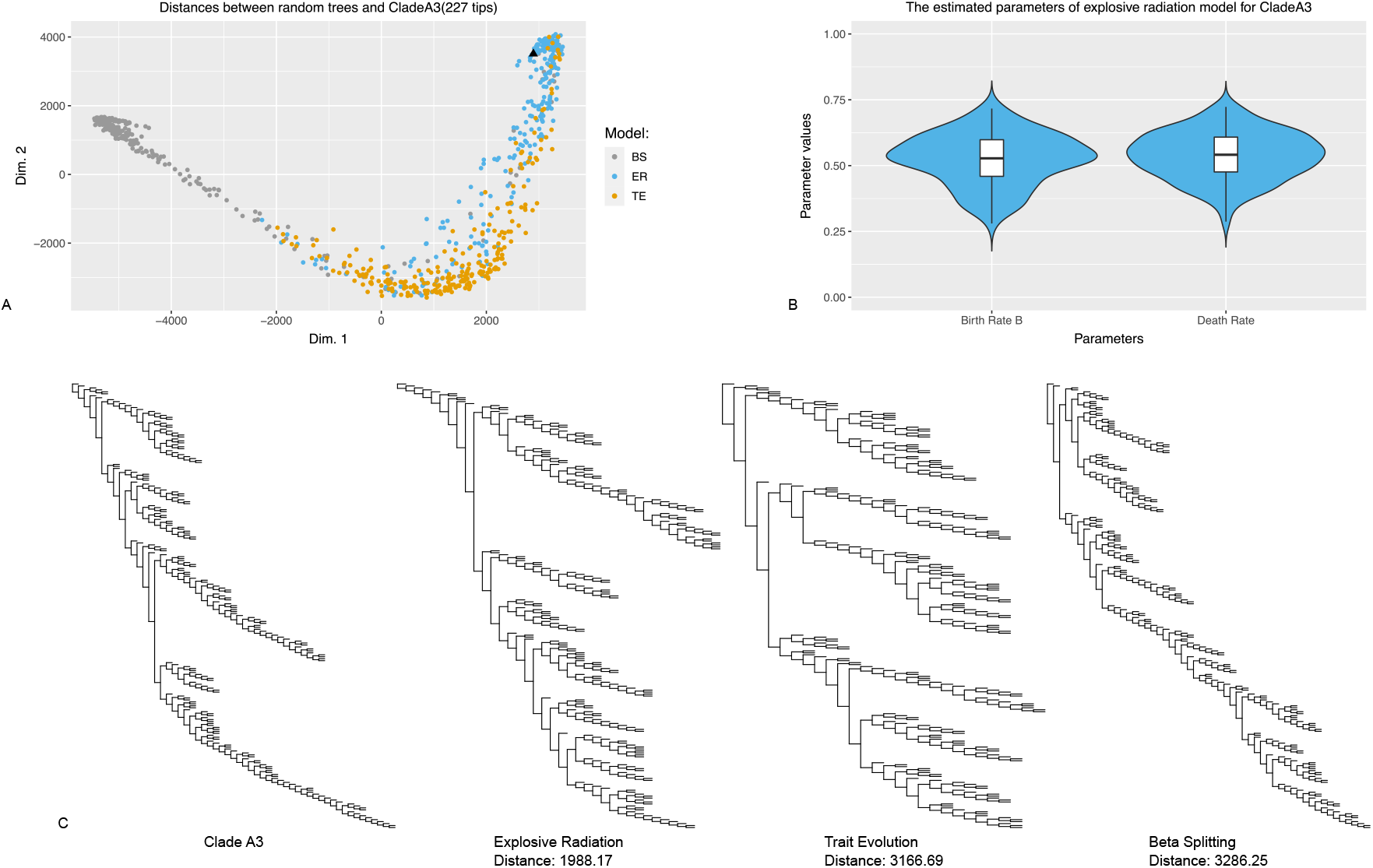
A: The MDS plots of the polynomial distances between the random trees generated by the three different models and influenza virus clade A3. B: the distribution of the estimated parameters of clade A3 over 100 replicates. C: the plots of clade A3 and the nearest random trees generated by the three different models.

Lastly, we plot, in Figure 5 C, clade A3 and the nearest random trees to the clade from each model in the sets of 250 random trees displayed in Figure 5 A. The polynomial distances between a pair of 227-tip trees has the upper bound of 12,882. Assuming the distribution of the pairwise distances will be normal as displayed in Supplementary Figure 1, all of the polynomial distances between clade A3 and the trees in Figure 5 C are below average pairwise distances of 227-tip trees. We also perform the same analysis on clade 3c3.B which has 117 tips (Supplementary Figure 7). Throughout our comparisons between simulated and real trees, we note that we have not simulated realistic total populations of either HIV or influenza in the relevant settings and then down-sampled to match the sizes of observed trees as this would be infeasible. This affects the interpretation of our estimates.

## Discussion

We have introduced a new way to describe and analyze phylogenetic trees using a polynomial that uniquely characterizes trees. We compare the polynomial to other indices and methods describing tree shapes. The polynomial is easy to compute and it has the advantage of describing trees in full resolution, that is, the descriptions are different if and only if the two tree shapes are not isomorphic. Moreover, the polynomials have the potential to be extended to record information about the branch lengths.

We also introduced some basic methods for tree analysis using the polynomial. The methods discussed in this paper include a tree metric, a parameter estimation method based on the metric, and a naive Bayes classifier directly trained by the coefficients of the polynomials. We chose these simple and tractable methods to show that the polynomial can be utilized in likelihood free methods for various tasks in analyzing phylogenetic trees. These polynomial based methods can distinguish trees from different models and different data sets, help estimate parameters, and aid in model selection. We have also applied these polynomial based methods to estimate parameters and select the best-fit model for the chosen WHO influenza virus clades. The results show that the tree shapes of the influenza clades are most similar to random trees generated by either the explosive radiation model or the trait evolution model, both of which are based on the birth-death process compared to the beta splitting model which is based on the Markov branching process. We also computed the nearest (in the polynomial distance) trees from each model to a WHO-defined influenza clade. This information, together with the distribution of the pairwise polynomial distances between trees being normal, can be used to assess how well a simulated tree resembles a tree reconstructed from data.

The simple methods used in this paper for parameter estimation and model selection can be improved in terms of computation efficiency among other aspects. And indeed, in estimation problems, it may be best to collect a wide range of tree descriptors (including polynomial coefficients, scalar summaries such as the Sackin and Colless imbalance measures and other high-resolution characterizations of the tree) (Saulnier, 2017), and let feature selection sort out which are best for a particular problem. Different models and data will yield trees with different features, and in some of these, simple scalar summary statistics may perform well. Our results show that in our simulation examples the polynomial coefficients are informative and would likely add to such an analysis, probably with the most benefit where scalar imbalance measures do not contain sufficient information about trees to perform the desired estimation task. Characterizing trees in the polynomial’s high-resolution metric way also allows selection of the closest tree to a tree from data, and visualizations of the space of trees derived from a model or datasets of interest. The polynomial can be used to obtain novel features or pseudo-metrics for clustering and estimation; as an example, the binary differences (Choi, 2010) can be used to improve clustering for the influenza and HIV trees (Supplementary Figure 6).

Our polynomial is not the only one that uniquely represents rooted binary trees. Other polynomials, such as the ones introduced in (Andrén, 2009), (Chaudhary, 1991), (Negami, 1996) and (Botti, 1993), (Matsen, 2012) are also good candidates for tree analysis. Thus it is worth investigating more about how these different polynomials can be used to analyze phylogenetic trees and how different results can they yield. The computation of most of these polynomials requires going through all subtrees or all permutations of a given size, which can be computationally heavy, while the polynomial used in this paper has a recursive formula which makes the computation more efficient.

To compare trees with different sizes is another challenge in tree comparison. In this paper, we have compared trees with the same number of tips and we have proposed a way to compare trees with different sizes. In our demonstration *treeverse*, trees with different sizes are compared and the distances between the trees are visualized by an interactive 3-D MDS plot. There are various ways to compare the coefficient vectors and compare trees with different sizes, but for trees whose sizes are drastically different, the sizes naturally remain a dominating factor in the resulting tree comparisons.

Because polynomial coefficients can be treated as vectors, and vectors give rise to metrics, there are alternative metrics that can be defined using tree polynomials (both those used here and others (Andrén, 2009; Chaudhary, 1991; Negami, 1996)). Once trees are encoded as vectors, a range of regression, inference and dimension reduction and other machine learning tools can, as a result, be applied to trees. In addition, other tree shape statistics or further information about the trees (including measures of branch length) can easily be appended to the vectors to integrate distinct sources of data. This provides a scheme to study phylogenetic trees comprehensively.

There remains considerable scope to improve the clustering and classification tools used here, which we used to demonstrate that parameter estimation and model choice can be done. One challenge in this work is that there are too many polynomial coefficients; however, feature selection, hyperparameter optimization and dimension reduction tools could be used to reduce the number of features in a systematic way. Furthermore, we used one- or two-dimensional estimation tasks as demonstrations. Realistic models of evolution are likely to contain multiple parameters (for example, time-dependent speciation and extinction rates; intra- and inter-group competition parameters, relative fitness), so more advanced and modern statistical inference tools could be considered. The simpler estimation we have provided is a proof of principle for using polynomial coefficients in estimation tasks.

## Acknowledgements

This work was supported by the grant of the Federal Government of Canada’s Canada 150 Research Chair program to Dr. Caroline Colijn. We would like to thank Art Poon, who provided the HIV trees.

## Supplementary Material

### The polynomial metric

We prove that the polynomial metric is a genuine metric. It is easy to check that *d*(*T*_1_, *T*_2_) = 0 if and only if *T*_1_ ≃ *T*_2_, and *d*(*T*_1_, *T*_2_) = *d*(*T*_2_, *T*_1_). We show that the triangular inequality is true for the metric, that is, *d*(*T*_1_, *T*_3_) ≤ *d*(*T*_1_, *T*_2_) + *d*(*T*_2_, *T*_3_). We only need to prove the following inequality holds for *a, b, c* ≥ 0.

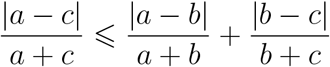

Note that if *a* ≥ *c* ≥ *b* or *c* ≥ *a* ≥ *b*, we have

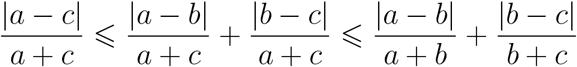

If *a* ≥ *b* ≥ *c*, we have

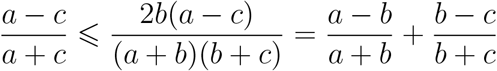

This is equivalent to *b*^2^ + *ac* ≤ *ab* + *bc*, which is true because *ac* − *bc* ≤ *ab* − *b*^2^. Similarly, the equality also holds when *c* ≥ *b* ≥ *a*.

If *b* ≥ *a* ≥ *c*, we have

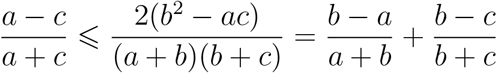

This is equivalent to *ab*(*b* − *a*) + 3*c*(*b*^2^ − *a*^2^) + *c*^2^(*b* − *a*) ≥ 0, which is true as *b* ≥ *a*. Similarly, the equality also holds when *b* ≥ *c* ≥ *a*. Therefore the polynomial metric is a genuine metric.

**Supplementary Figure 1.**
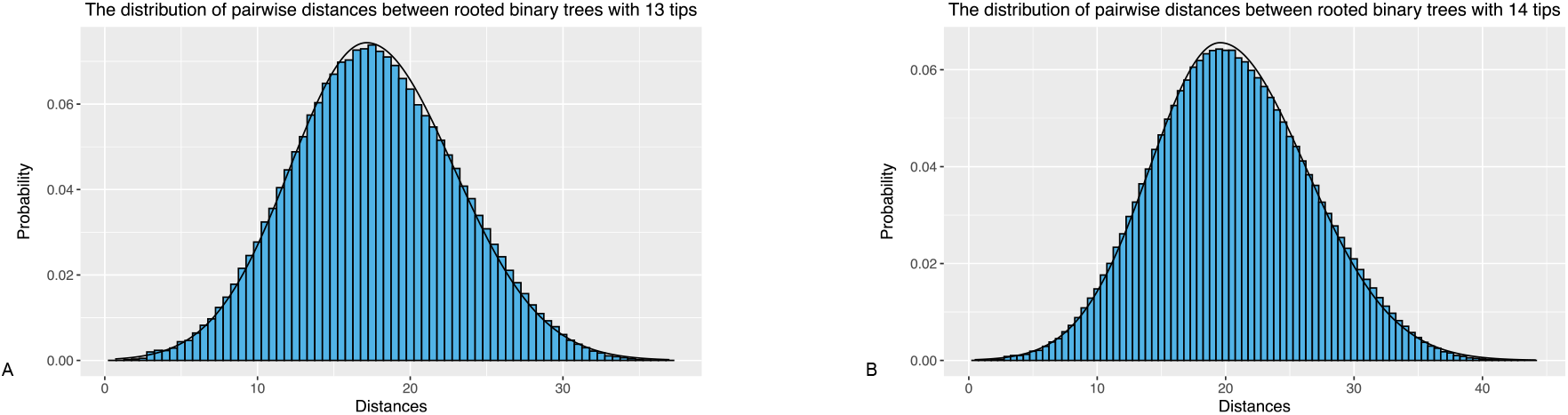
A-B: the distribution of all pairwise polynomial distances between all rooted binary trees with 13 and 14 tips, where the black solid curves are normal fits.

The distribution of polynomial distances between all pairs of trees with *n* tips resembles a normal distribution. Supplementary Figure 1 displays the distribution for trees with 13 and 14 tips, where the black solid curves are normal fits. For the distribution for 13-tip trees, the estimated mean value is 17.70, the estimated standard deviation is 5.37, and Shapiro-Wilk normality test has W of 0.99 and p-value of 6.21 × 10^−15^. For the distribution for 14-tip trees, the estimated mean value is 20.54, the estimated standard deviation is 6.10, and Shapiro-Wilk normality test has W of 0.99 and p-value of 4.43 × 10^−15^.

### Parameter estimation and model selection

We show the supplementary results of parameter estimation and model selection in complement to the figures displayed in the main result section. Supplementary Figure 2 shows the results of parameter estimation for 750-tip beta splitting trees, 250-tip and 500-tip explosive radiation trees. Supplementary Table 1, 2 and 3 show the summaries of the estimation. Supplementary Figure 3 shows the results of model selection for 250-tip and 750-tip random trees generated by the three models. Supplementary Figure 4 shows the results parallel to the results displayed in Figure 3 with subtree size distributions instead of polynomials.

**Supplementary Table 1.**
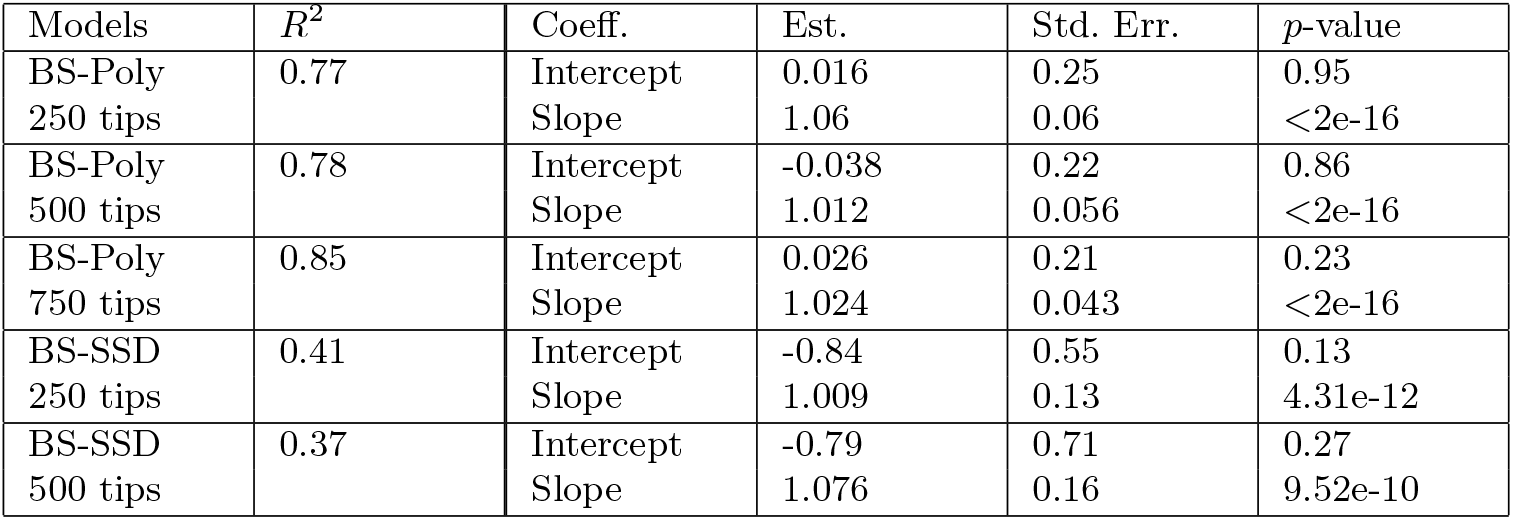
The summary of linear fit of the real parameter and the estimated parameter for beta splitting trees.

| Models | $R^2$ | Coeff. | Est. | Std. Err. | $p$-value |
| --- | --- | --- | --- | --- | --- |
| BS-Poly<br>250 tips | 0.77 | Intercept<br>Slope | 0.016<br>1.06 | 0.25<br>0.06 | 0.95<br><2e-16 |
| BS-Poly<br>500 tips | 0.78 | Intercept<br>Slope | -0.038<br>1.012 | 0.22<br>0.056 | 0.86<br><2e-16 |
| BS-Poly<br>750 tips | 0.85 | Intercept<br>Slope | 0.026<br>1.024 | 0.21<br>0.043 | 0.23<br><2e-16 |
| BS-SSD<br>250 tips | 0.41 | Intercept<br>Slope | -0.84<br>1.009 | 0.55<br>0.13 | 0.13<br>4.31e-12 |
| BS-SSD<br>500 tips | 0.37 | Intercept<br>Slope | -0.79<br>1.076 | 0.71<br>0.16 | 0.27<br>9.52e-10 |

**Supplementary Table 2.** The summary of linear fit of the real parameter and the estimated parameter λ_B_ for explosive radiation trees.

| Models | $R^2$ | Coeff. | Est. | Std. Err. | $p$-value |
| --- | --- | --- | --- | --- | --- |
| ER-Poly<br>250 tips | 0.035 | Intercept<br>Slope | 0.33<br>0.46 | 0.15<br>0.25 | 0.033<br>0.063 |
| ER-Poly<br>500 tips | 0.043 | Intercept<br>Slope | 0.36<br>0.43 | 0.13<br>0.21 | 0.006<br>0.037 |
| ER-Poly<br>750 tips | 0.09 | Intercept<br>Slope | 0.29<br>0.58 | 0.14<br>0.22 | 0.048<br>0.011 |
| ER-SSD<br>750 tips | 0.07 | Intercept<br>Slope | 0.40<br>0.44 | 0.10<br>0.16 | 0.001<br>0.008 |

**Supplementary Table 3.** The summary of linear fit of the real parameter and the estimated parameter *μ* for explosive radiation trees.

| Models | $R^2$ | Coeff. | Est. | Std. Err. | $p$-value |
| --- | --- | --- | --- | --- | --- |
| ER-Poly<br>250 tips | 0.24 | Intercept<br>Slope | 0.12<br>0.67 | 0.05<br>0.12 | 0.016<br>1.87e-07 |
| ER-Poly<br>500 tips | 0.46 | Intercept<br>Slope | 0.067<br>0.84 | 0.041<br>0.091 | 0.11<br>7.44e-15 |
| ER-Poly<br>750 tips | 0.69 | Intercept<br>Slope | 0.034<br>0.94 | 0.031<br>0.075 | 0.28<br><2e-16 |
| ER-SSD<br>750 tips | 0.59 | Intercept<br>Slope | -0.008<br>1.40 | 0.035<br>0.12 | 0.815<br><2e-16 |

In Supplementary Figure 5, we show the importance of features in the naive Bayes classifiers used for model selection with both subtree size distributions and polynomials. As the naive Bayes classifiers assume independence of variables, the Shannon entropy reflects the importance of the features, where a feature with smaller entropy means the feature is more important in the classifier.

**Supplementary Figure 2.**
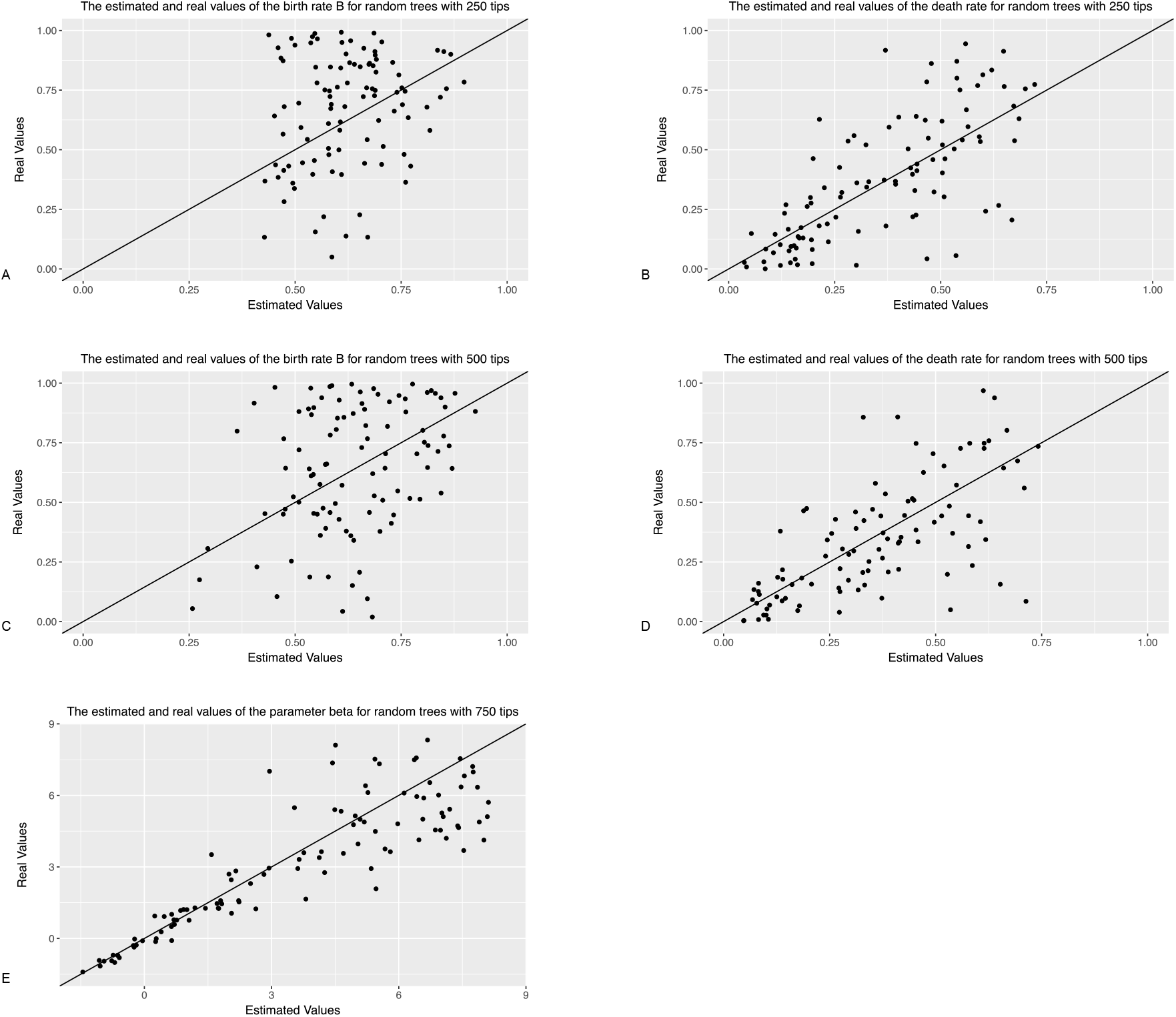
A-B: the comparisons between the real parameters and the estimated parameters of the explosive radiation random trees with 250 tips using polynomials. C-D: the comparisons between the real parameters and the estimated parameters of the explosive radiation random trees with 500 tips using polynomials. E: the comparisons between the real parameter and the estimated parameter of the beta splitting random trees with 750 using polynomials.

### Polynomial binary differences

Binary differences, based on presence and absence of components, though in general not metrics, are one of the commonly used indices in, for example, taxonomic, ecologic, biogeographic comparison and classification (Choi, 2010). They provide effective insights about clusters though they are not metrics in general. We define the polynomial binary differences used in this paper by the number of terms that are present in the polynomial of one tree but are absent in the polynomial of the other. More precisely, the binary difference of two trees *T*_1_ and *T*_2_ are calculated by counting the number of terms that are present in *P*(*T*_1_, *x, y*) but are absent in *P*(*T*_2_, *x, y*), or the number of terms that are present in *P*(*T*_2_, *x, y*) but are absent in *P*(*T*_1_, *x, y*). This provides another way to compare polynomials (trees). Supplementary Figure 6 shows the results of *k*-medoids clustering on the binary differences of the influenza trees and the HIV trees, which are better than the polynomial metric in this task.

**Supplementary Figure 3.**
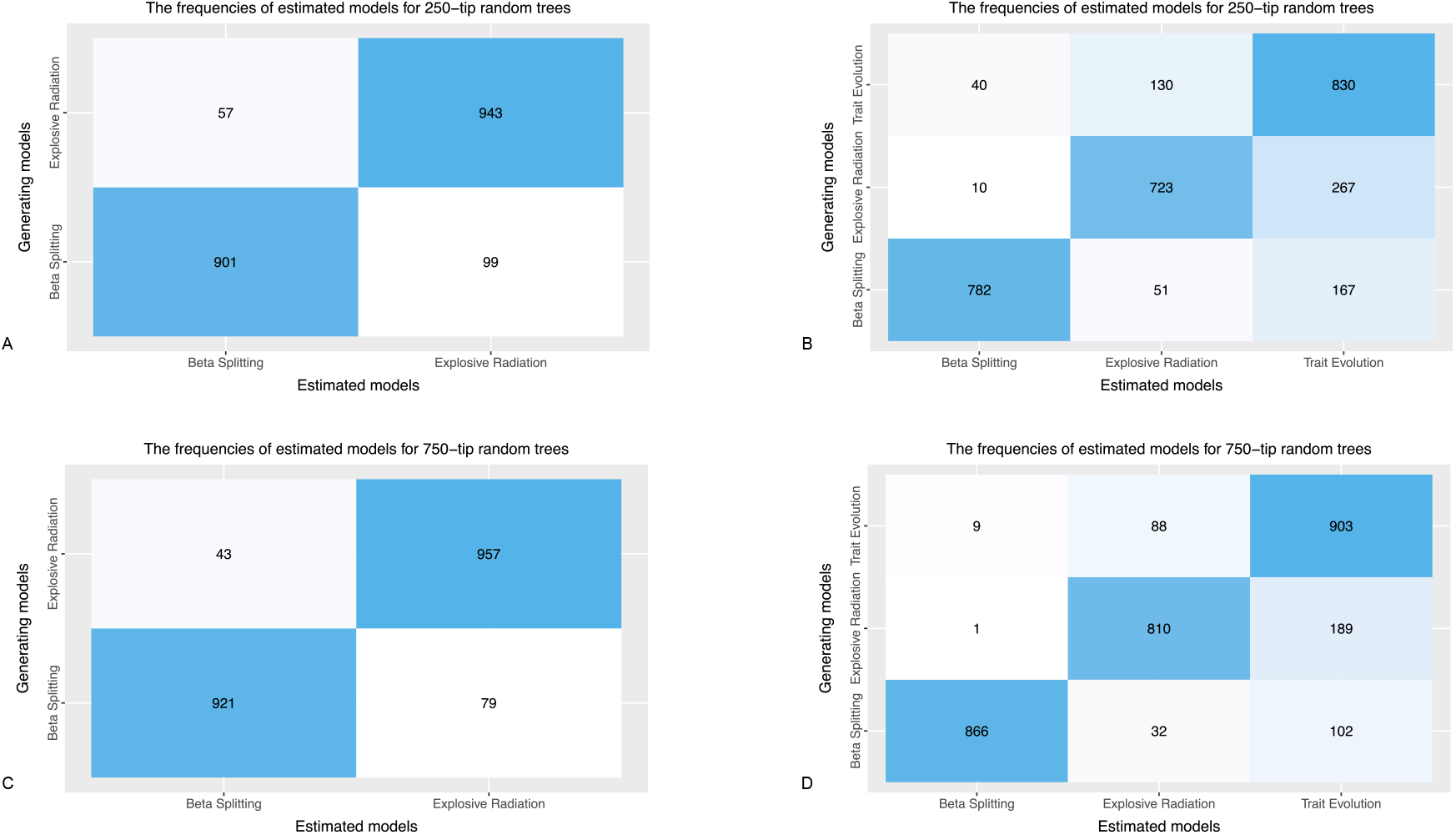
A-B: the results of using naive Bayes classifiers to select the model generating random trees with 250 tips using polynomials. C-D: the results of using naive Bayes classifiers to select the model generating random trees with 750 tips using polynomials.

### WHO influenza clades

For clade 3c3.B, the 95% confidence interval of the birth rate λ_*B*_ is (0.56, 0.60) and the 95% confidence interval of the death rate *μ* is (0.58, 0.62). The 95% confidence interval of *R*_0_ of the clade is (0.918, 1.013).

**Supplementary Figure 4.**
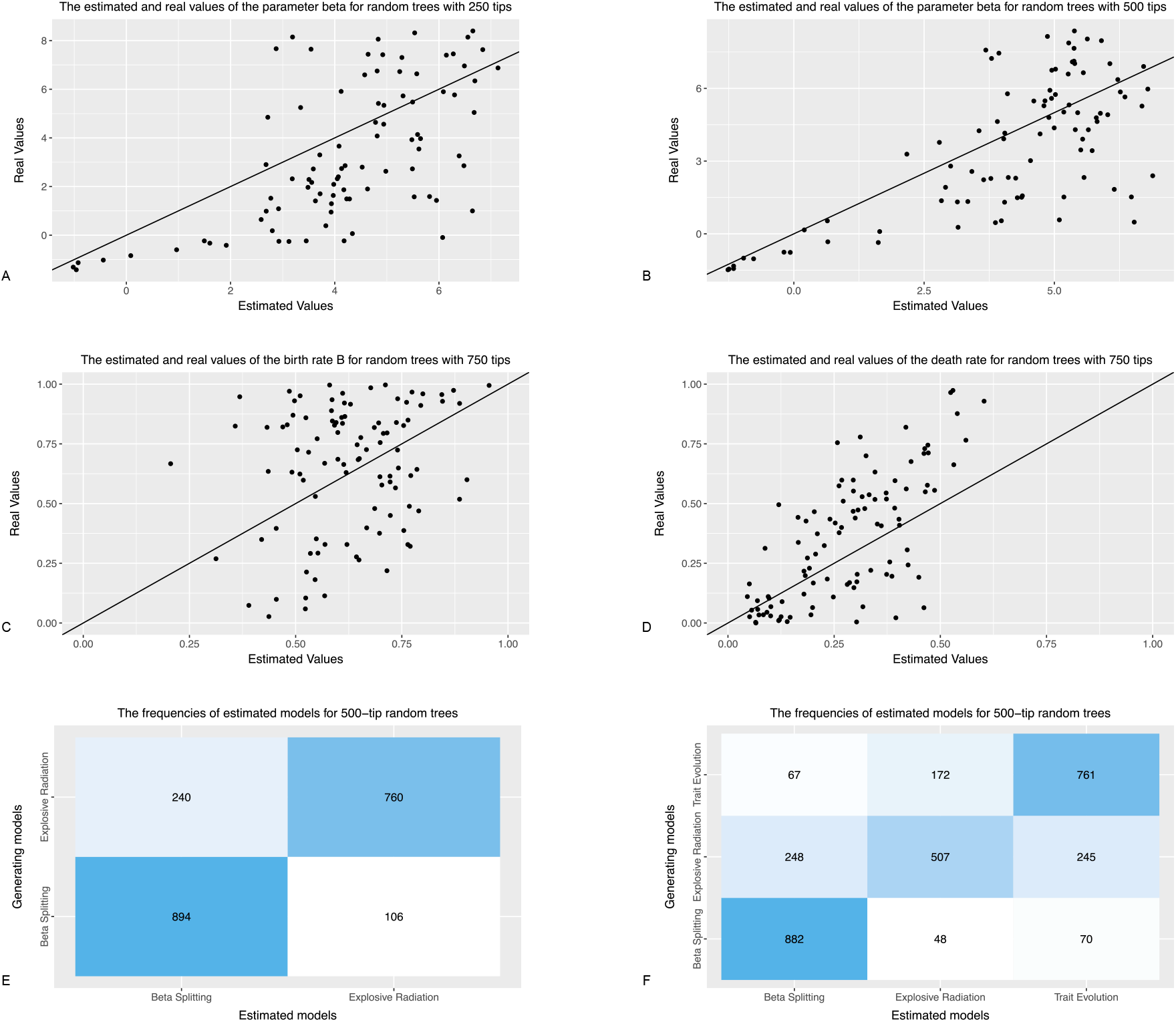
A-B: the comparisons between the real parameter and the estimated parameter of the beta splitting random trees with 250 tips and 500 tips using subtree size distributions. C-D: the comparisons between the real parameters and the estimated parameters of the explosive radiation random trees with 750 tips using subtree size distributions. E-F: the results of using naive Bayes classifiers to select the model generating random trees with 500 tips using subtree size distributions.

**Supplementary Figure 5.**
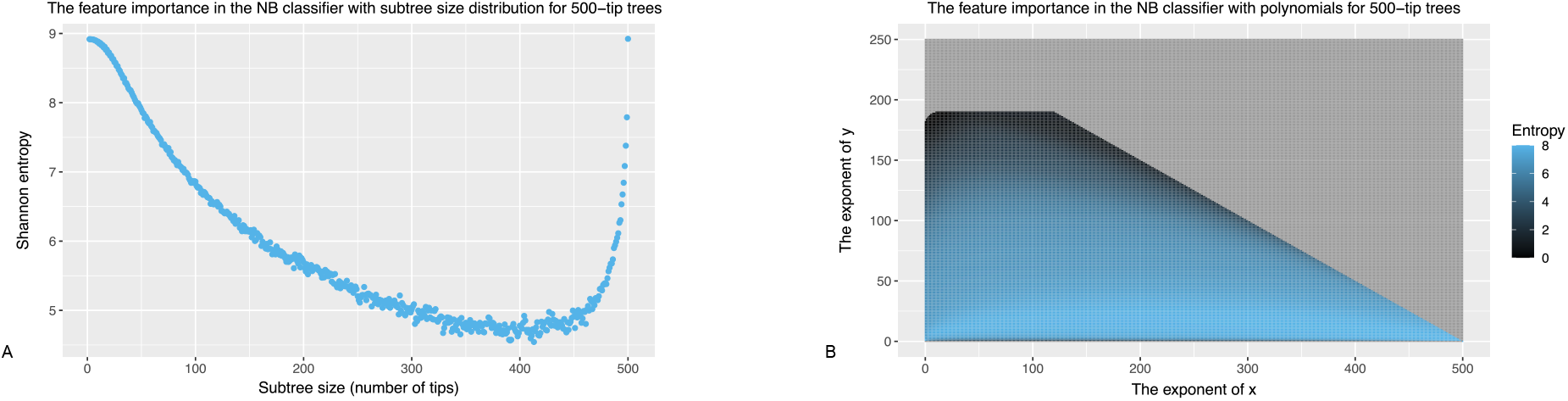
A: the feature importance (Shannon entropy) in the naive Bayes classifier used for model selection with subtree size distributions. B: the feature importance (Shannon entropy) in the naive Bayes classifier used for model selection with polynomials.

**Supplementary Figure 6.**
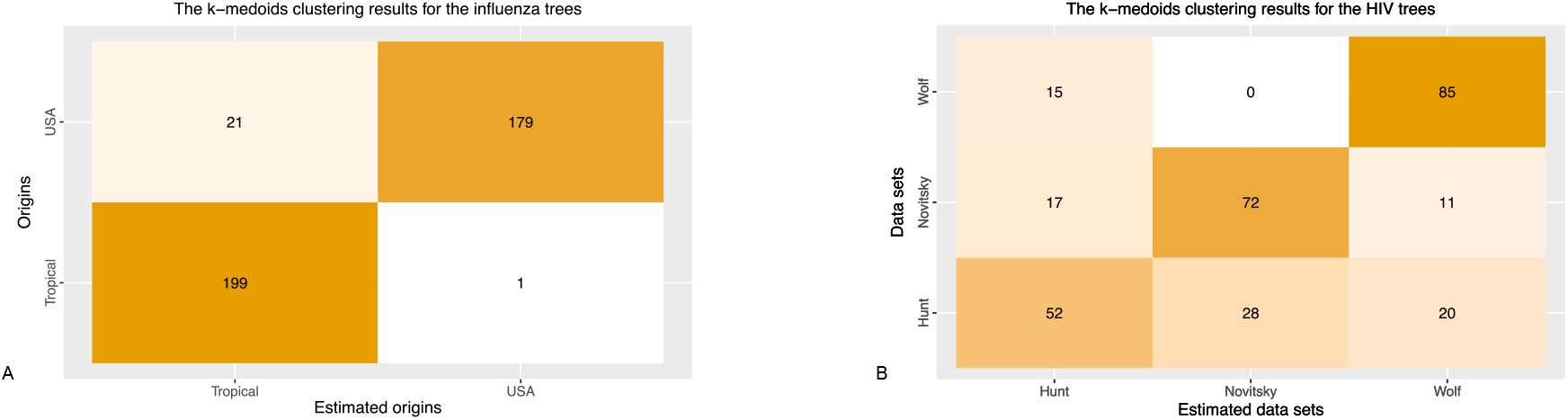
A: the results of *k*-medoids clustering for the influenza trees using the polynomial binary differences. B: the results of *k*-medoids clustering for the HIV trees using the polynomial binary differences.

**Supplementary Figure 7.**
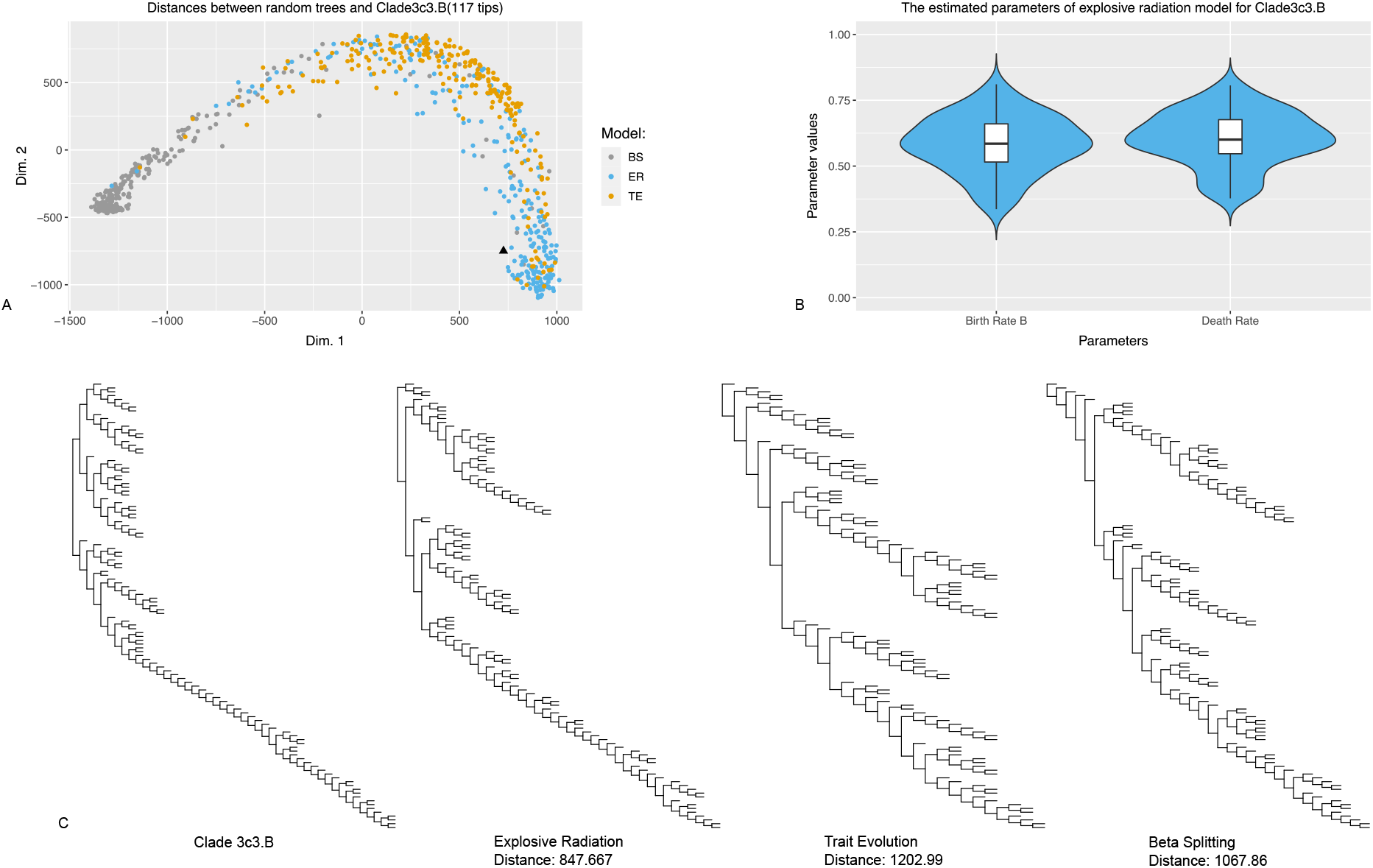
A: the MDS plots of the polynomial distances between the random trees generated by the three different models and the clade 3c3.B. B: the distribution of the estimated parameters of the clade 3c3.B over 100 replicates. C: the plots of the clade A3 and the nearest random trees generated by the three different models.

## Notes

### Competing Interest Statement

The authors have declared no competing interest.

